# Neuropeptide VF neurons promote sleep via the serotonergic raphe

**DOI:** 10.1101/2019.12.27.889402

**Authors:** Daniel A. Lee, Grigorios Oikonomou, Tasha Cammidge, Young Hong, David A. Prober

## Abstract

Although several sleep-regulating neurons have been identified, little is known about how they interact with each other for sleep/wake control. We previously identified neuropeptide VF (NPVF) and the hypothalamic neurons that produce it as a sleep-promoting system (Lee et al., 2017). Here we use zebrafish to describe a neural circuit in which *neuropeptide VF* (*npvf*)-expressing neurons control sleep via the serotonergic raphe nuclei (RN), a hindbrain structure that promotes sleep in both diurnal zebrafish and nocturnal mice. Using genetic labeling and calcium imaging, we show that *npvf*-expressing neurons innervate and activate serotonergic RN neurons. We additionally demonstrate that optogenetic stimulation of *npvf-*expressing neurons induces sleep in a manner that requires NPVF and is abolished when the RN are ablated or lack serotonin. Finally, genetic epistasis demonstrates that NPVF acts upstream of serotonin in the RN to maintain normal sleep levels. These findings reveal a novel hypothalamic-hindbrain circuit for sleep/wake control.

## INTRODUCTION

While several sleep- and wake-promoting neuronal populations have been identified (reviewed in (Bringmann, 2018; Liu and Dan, 2019; Saper and Fuller, 2017; Scammell et al., 2017)), characterizing and understanding the functional and hierarchical relationships between these populations is essential for understanding how the brain regulates sleep and wake states (Oikonomou and Prober, 2017). Recent evidence from zebrafish and mice demonstrate that the serotonergic raphe nuclei (RN) are critical for the initiation and maintenance of sleep (Oikonomou et al., 2019), in contrast with previous models suggesting a wake-promoting role for the RN that were largely based on their wake-active nature (Saper and Fuller, 2017; Scammell et al., 2017; Weber and Dan, 2016). In zebrafish, mutation of *tryptophan hydroxylase 2* (*tph2*), which is required for serotonin (5-HT) synthesis in the RN, results in reduced sleep, sleep depth, and homeostatic response to sleep deprivation (Oikonomou et al., 2019). Pharmacological inhibition of 5-HT synthesis or ablation of the RN also results in reduced sleep. Consistent with a sleep-promoting role for the raphe, optogenetic stimulation of raphe neurons results in increased sleep. Similarly, in mice, ablation of the RN results in increased wakefulness and an impaired homeostatic response to sleep deprivation, whereas tonic optogenetic stimulation of the RN at a rate similar to their baseline pattern of activity induces NREM sleep. These complementary results in zebrafish and mice (Oikonomou et al., 2019), along with classical ablation and pharmacological studies (Ursin, 2008), indicate an evolutionarily conserved role for the serotonergic system in promoting vertebrate sleep. However, it is unclear how the RN are themselves regulated to promote sleep.

Trans-synaptic retrograde viral tracing studies have identified substantial inputs to the RN from hypothalamic neurons in the lateral hypothalamic area, tuberomammillary nucleus, and dorsomedial nucleus, regions implicated in sleep-wake regulation (Pollak Dorocic et al., 2014; Ren et al., 2018; Weissbourd et al., 2014). However, it is unknown whether any of these or other populations act upon the RN to promote sleep. One candidate neuronal population expresses the sleep-promoting neuropeptide VF (NPVF) in ∼25 neurons in the larval zebrafish hypothalamus (Lee et al., 2017). Overexpression of *npvf* or stimulation of *npvf*-expressing neurons results in increased sleep, whereas pharmacological inhibition of NPVF signaling or ablation of *npvf*-expressing neurons results in reduced sleep (Lee et al., 2017). While it is unknown how the NPVF system promotes sleep, these neurons densely innervate a region of the hindbrain that is consistent with the location of the RN (Lee et al., 2017; Madelaine et al., 2017). Because perturbations of the NPVF system and RN have similar effects on sleep, and processes of *npvf*-expressing neurons are found near the RN, we hypothesized that the NPVF system promotes sleep via the RN. To test this hypothesis, we used genetics, optogenetics, and chemogenetics to dissect the functional relationship between these two neuronal populations. Our results support the hypothesis that the NPVF system promotes sleep via the RN, thus revealing a novel hypothalamus-hindbrain neural circuit for sleep-wake control.

## RESULTS

### NPVF neurons densely innervate the serotonergic inferior raphe nuclei

In most vertebrates, the RN are the main source of serotonergic innervation in the brain. In mammals, the RN are divided into a rostral or superior group that lies on the midbrain/pons boundary (groups B5–B9) (Dahlstroem and Fuxe, 1964), and a caudal or inferior group in the medulla (groups B1–B3). Similarly, in zebrafish larvae, the RN are divided into the superior raphe (SRa) and inferior raphe (IRa) (Lillesaar et al., 2009).

To explore whether the NPVF system may promote sleep via the RN, we first performed a detailed histological analysis of these populations using *Tg(npvf:eGFP)* (Lee et al., 2017) and *Tg(npvf:KalTA4); Tg(UAS:nfsb-mCherry)* (Agetsuma et al., 2010; Lee et al., 2017) animals, which specifically label *npvf*-expressing neurons. As previously described (Lee et al., 2017), the soma of *npvf*-expressing neurons are present exclusively in the dorsomedial hypothalamus at 6 days post-fertilization (dpf) (**Figure 1A,B,D**). These neurons send dense and local ramifying projections into the hypothalamus (**Figure 1B,D**), as well as longer-range projections into the telencephalon and hindbrain, with a prominent convergence of these projections at the rostral IRa, as confirmed using 5-HT immunohistochemistry (IHC) (**Figure 1B-1K, Figure 1 – figure supplement 1A-B**). These projections form a dense bundle just ventral to the soma of the IRa and also extend dorsally where they appear to make multiple contacts with IRa somas. As additional confirmation of this interaction, we mated *Tg(npvf:KalTA4); Tg(UAS:nfsb-mCherry)* (Agetsuma et al., 2010; Lee et al., 2017) animals, in which NPVF neurons and their processes are labeled with mCherry, to *Tg(tph2:eNTR-mYFP)* animals, in which the SRa and IRa are labeled with membrane-targeted YFP (Oikonomou et al., 2019). We observed apparent direct contacts of NPVF neuron fibers with mYFP-labeled IRa soma and fibers (**Figure 1 – figure supplement 1C-E**), consistent with a direct interaction between NPVF and IRa neurons.

**Figure 1.**
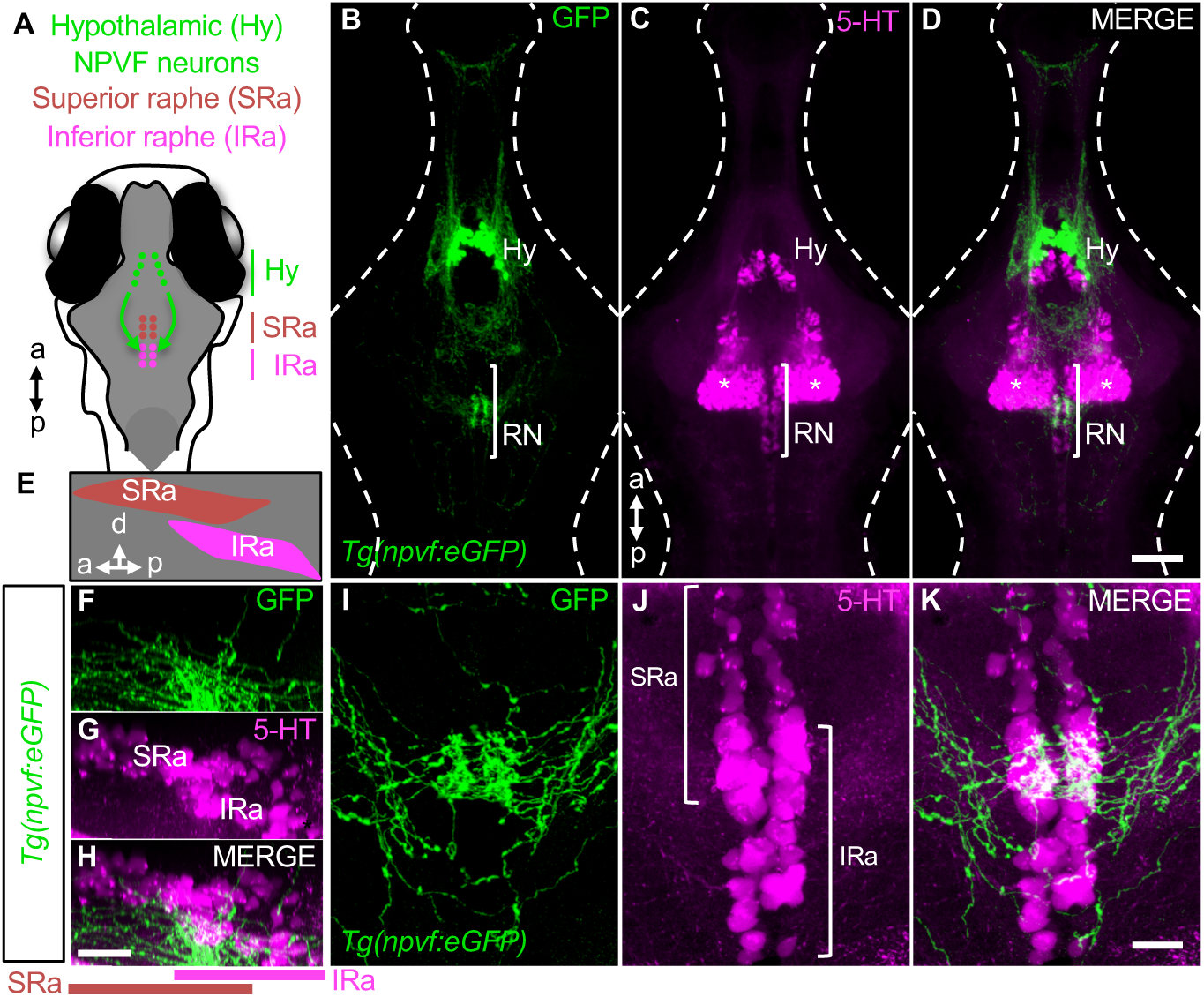
Hypothalamic NPVF neurons project to the serotonergic IRa. (**A,E**) Schematic 6-dpf zebrafish brain showing location of hypothalamic (Hy) NPVF neurons (green), and the serotonergic SRa (red) and IRa (magenta). a, anterior; p, posterior; d, dorsal. (**B-D**) Maximum intensity projection of a brain from a 6-dpf *Tg(npvf:eGFP)* animal (78 μm thick). *npvf*-expressing neurons in the hypothalamus project to the serotonergic raphe nuclei (RN) in the hindbrain (bracket). 5-HT immunohistochemistry labels the RN (bracket), as well as serotonergic populations in the ventral hypothalamus (asterisks) and pretectum. The bracketed region in (**B-D**) is shown at higher magnification in (**I-K**) as a maximum intensity projection (50.5 μm thick), with a sagittal view shown in (**E-H**). Single optical sections are shown in **Figure S1**. Scale: 50 μm (**B-D**), 20 μm (**F-H)** and 10 μm (**I-K**).

### Optogenetic stimulation of NPVF neurons results in activation of serotonergic IRa neurons

Based on our histological observations, we hypothesized that NPVF neurons are functionally connected to serotonergic IRa neurons, and that stimulation of NPVF neurons should thus activate IRa neurons. To test this hypothesis, we used *Tg(npvf:ReaChR-mCitrine);Tg(tph2:GCaMP6s-tdTomato)* animals (Lee et al., 2017; Oikonomou et al., 2019) to optogenetically stimulate NPVF neurons while monitoring the activity of IRa neurons. Because neurons in the RN are responsive to visible light (Cheng et al., 2016), we used invisible 920 nm two-photon light at low laser power to excite GCaMP6s fluorescence. We also used 920 nm two-photon light, applied at higher power, to stimulate ReaChR in NPVF neurons.

To verify that this stimulation paradigm indeed results in stimulation of NPVF neurons, we first tested *Tg(npvf:ReaChR-mCitrine)*; *Tg(npvf:GCaMP6s-tdTomato)* animals (Lee et al., 2017; Lee et al., 2019). We first recorded baseline GCaMP6s fluorescence in *npvf*-expressing neurons, then optogenetically stimulated these neurons, and then recorded post-stimulation GCaMP6s fluorescence in these neurons (**Figure 2 – figure supplement 1**). In the frame after compared to the frame before optogenetic stimulation, we observed a 165% increase in GCaMP6s fluorescence in NPVF neurons in *Tg(npvf:ReaChR-mCitrine)* animals (p<0.0005, Mann-Whitney test, 5 animals, 86 neurons) (**Figure 2 – figure supplement 1H,I**). In contrast, there was a 3% decrease in GCaMP6s fluorescence in the first frame after compared to the frame before optogenetic stimulation using the same imaging paradigm in *Tg(npvf:eGFP)* control animals (p<0.01, Mann-Whitney test, 5 animals, 95 neurons) (**Figure 2 – figure supplement 1I**). Thus, our stimulation paradigm results in robust ReaChR-dependent activation of *npvf*-expressing neurons.

**Figure 2.**
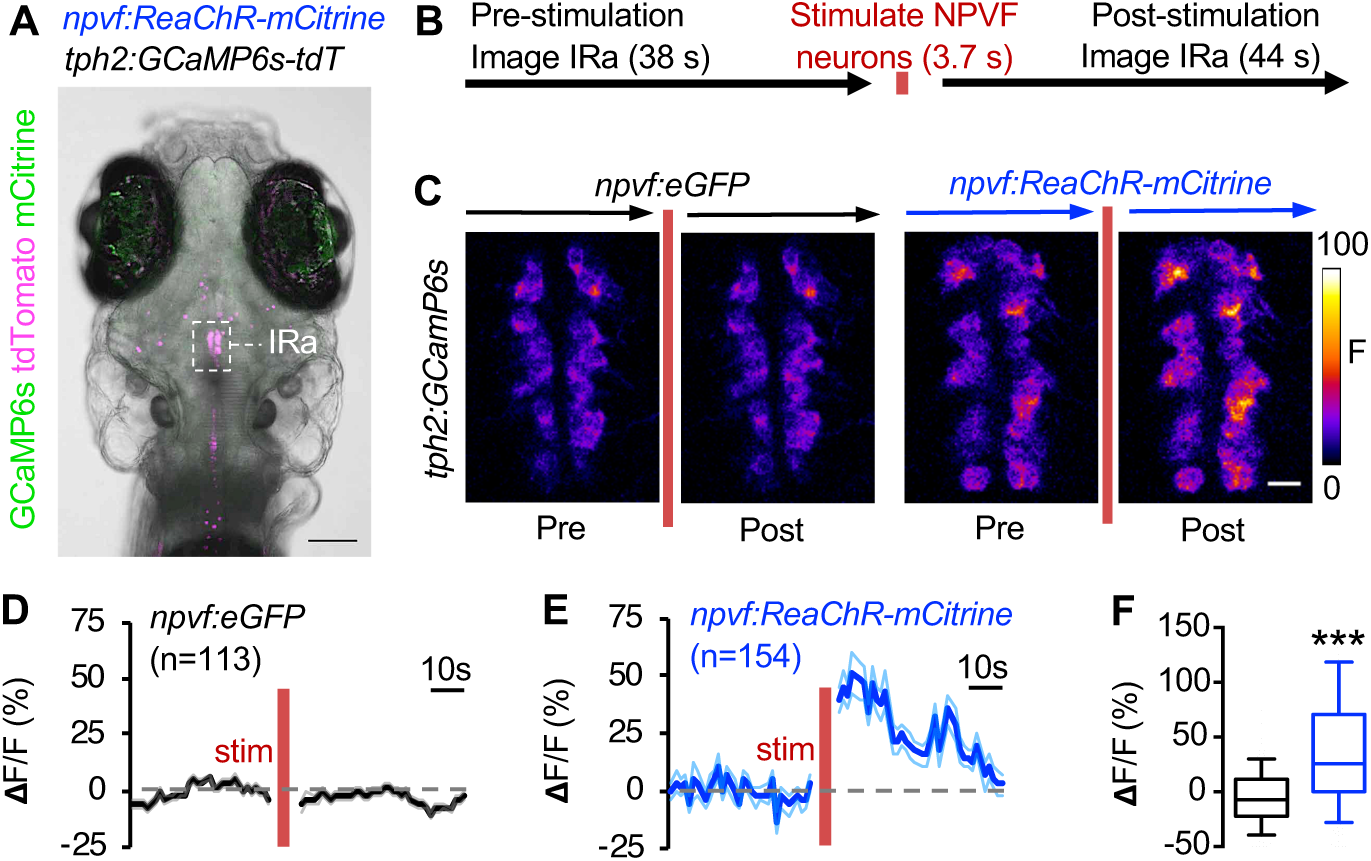
Optogenetic stimulation of NPVF neurons activates serotonergic IRa neurons. (**A-F**) 6-dpf *Tg(npvf:ReaChR-mCitrine); Tg(tph2:GCaMP6s-tdTomato)* and *Tg(npvf:eGFP); Tg(tph2:GCaMP6s-tdTomato)* animals are analyzed for GCaMP6s fluorescence levels in IRa neurons before and after optogenetic stimulation of NPVF neurons. (**A**) Example of a 6-dpf *Tg(npvf:ReaChR-mCitrine); Tg(tph2:GCaMP6s-tdTomato)* animal showing GCaMP6s and tdTomato fluorescence in IRa neurons. White box indicates the region of the IRa analyzed in subsequent panels. Green and magenta signals in the eyes are due to autofluorescence. (**B**) Baseline GCaMP6s fluorescence in the IRa was first recorded using two-photon 920 nm light at 8 mW laser power for 38 s (30 frames). NPVF neurons were then optogentically stimulated using 920 nm light at 38 mW at 2.7 Hz for 3.7 s. GCaMP6s fluorescence in the IRa was then immediately imaged again using 8 mW laser power for 44 s (35 frames). (**C**) Average GCaMP6s fluorescence in IRa neurons for 10 frames before (Pre) and after (Post) optogenetic stimulation of NPVF neurons. (**D,E**) Mean ± SEM GCaMP6s fluorescence of IRa neurons plotted as a function of time, showing pre-stimulation and post-stimulation evoked responses for *Tg(tph2:GCaMP6s-tdTomato)* animals that also contain a *Tg(npvf:eGFP)* (**D,F**) or *Tg(npvf:ReaChR-mCitrine)* (**E,F**) transgene. Dashed grey line indicates 0% **Δ**F/F. n=number of neurons quantified from 4 animals of each genotype. (**F**) Box plots quantify GCaMP6s **Δ**F/F values one frame after stimulation. ***p<0.0005, Mann-Whitney test. Scale: 100 μm (**A**), 10 μm (**C**).

We next used the same stimulation and imaging paradigm to ask whether optogenetic stimulation of NPVF neurons results in activation of neurons in the IRa using *Tg(npvf:ReaChR-mCitrine); Tg(tph2:GCaMP6s-tdTomato)* animals (Lee et al., 2017; Oikonomou et al., 2019). To do so, we first recorded baseline GCaMP6s fluorescence in *tph2*-expressing IRa neurons, then stimulated *npvf-*expressing neurons as described above, and then recorded post-stimulation GCaMP6s fluorescence in IRa neurons (**Figure 2A,B**). In the frame immediately following optogenetic stimulation, we observed a 40% increase in GCaMP6s fluorescence in IRa neurons (p<0.0001, Mann-Whitney test, 4 animals, 154 neurons) that gradually declined to baseline after ∼ 40 seconds (**Figure 2C, E,F**). This extended decay of GCaMP6s fluorescence is consistent with the prolonged effect expected for neuropeptide/G-protein coupled receptor (GPCR) signaling (van den Pol, 2012). Notably, while 74% of *tph2*-expressing neurons displayed an increase in GCaMP6s fluorescence immediately following stimulation (average ± SEM **Δ**F/F = 63±6%, 116 of 154 neurons), 26% of IRa neurons showed decreased fluorescence (average ± SEM **Δ**F/F = – 27±4%, 38 of 154 neurons), demonstrating a heterogeneity in response profiles among Ira neurons. These effects were due to optogenetic stimulation of NPVF neurons, as they were not observed in *Tg(npvf:eGFP)*; *Tg(tph2:GCaMP6s-tdTomato)* control animals (**Figure 2C,D,F**). These results demonstrate that stimulation of NPVF neurons results in activation of serotonergic IRa neurons, likely via neuropeptide/GPCR signaling.

### Loss of *npvf* does not enhance the *tph2* mutant sleep phenotype

The NPVF prepro-peptide is processed to produce three mature neuropeptides, RFRP 1-3 (Hinuma et al., 2000). We previously generated zebrafish that contain a frameshift mutation within the *npvf* gene that is predicted to encode a protein that contains RFRP1 but lacks RFRP2 and RFRP3 (Lee et al., 2017). We have shown that loss of NPVF signaling due to this mutation (Lee et al., 2017), or loss of 5-HT production in the RN due to mutation of *tph2* (Oikonomou et al., 2019), results in decreased sleep. Based on our observations that NPVF neurons project to and can stimulate serotonergic IRa neurons (**Figure 1**,**2**), we next tested the hypothesis that *npvf* and *tph2* act in the same genetic pathway to promote sleep. We tested this hypothesis by comparing the sleep of *npvf* −/−; *tph2* −/− animals to their heterozygous mutant sibling controls (**Figure 3**). We reasoned that if *npvf* and *tph2* promote sleep via independent genetic pathways, then animals lacking both genes should sleep more than either single mutant. In contrast, if *npvf* and *tph2* promote sleep in the same pathway, then loss of both genes should not result in an additive sleep phenotype. Similar to previous results, animals containing a homozygous mutation in either *npvf* or *tph2* slept less than heterozygous mutant sibling controls, with *tph2* mutants showing a stronger phenotype (**Figure 3B,C,E**). However, *npvf* −/−; *tph2* −/− animals did not sleep significantly more than their *npvf* +/−; *tph2* −/− siblings (**Figure 3D,E**). Thus, loss of *npvf* does not enhance the *tph2* mutant phenotype, consistent with the hypothesis that *tph2* acts downstream of *npvf* to promote sleep.

**Figure 3.**
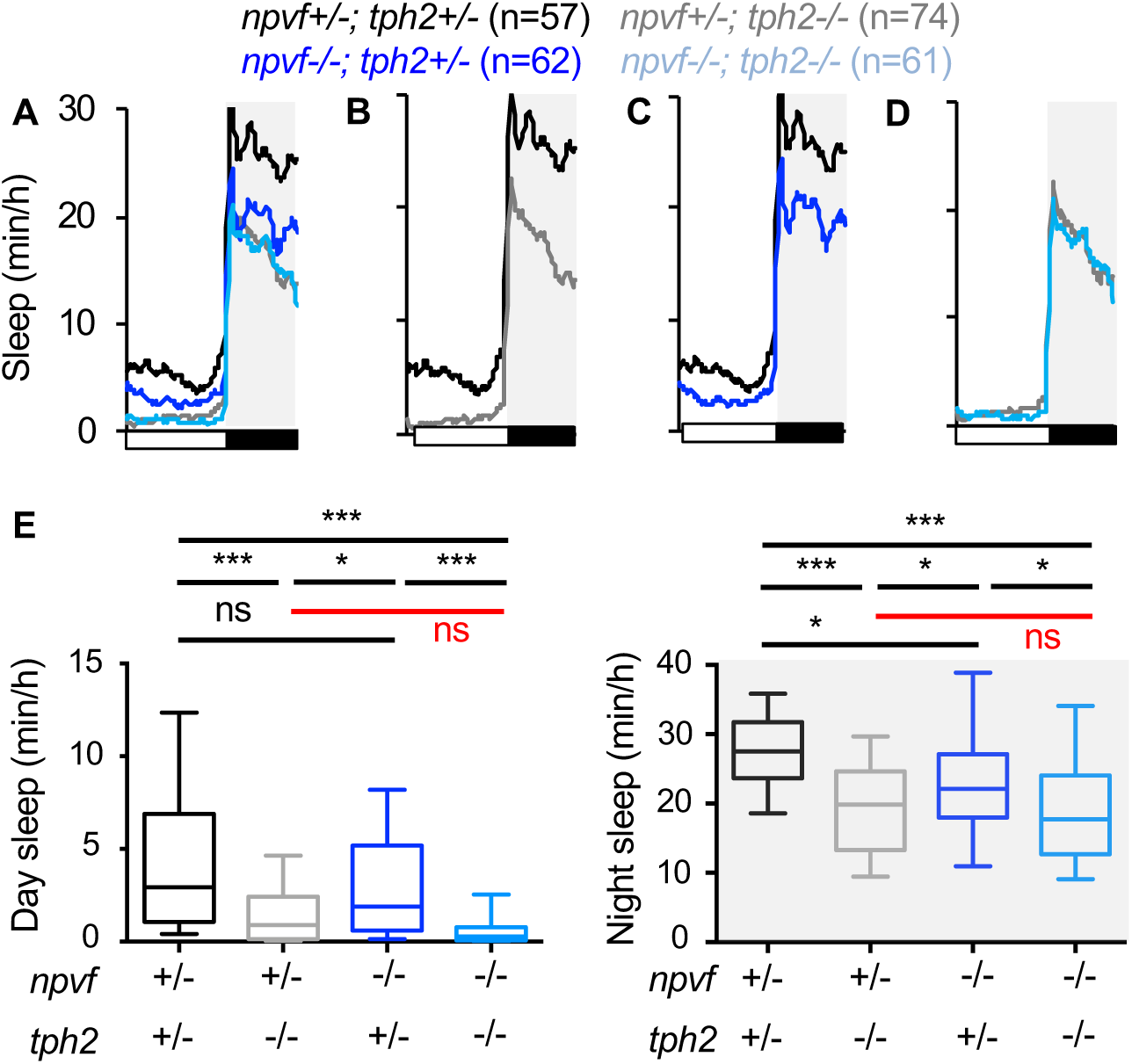
Loss of *npvf* does not enhance the *tph2* mutant sleep phenotype. (**A-D**) Average sleep for *npvf* +/−; *tph2* +/− (black), *npvf* −/−; *tph2* +/− (dark blue), *npvf* +/−; *tph2* −/− (gray), and *npvf* −/−; *tph2* −/− (light blue) siblings. The graph on the left shows data for all four genotypes. The three other graphs show the same data as separate pair-wise comparisons. (**E**) Box plots quantify sleep during the day (left) and night (right). White boxes indicate day. Black boxes and gray shading indicate night. n=number of animals. ns p>0.05, *p<0.05, **p<0.01, ***p<0.005, Two-way ANOVA with Holm-Sidak’s multiple comparisons test for each indicated pair-wise comparison. The comparison in red indicates no significant difference between *npvf* +/−; *tph2* −/− and *npvf* −/−; *tph2* −/− siblings.

### Sleep induced by stimulation of *npvf*-expressing neurons requires *npvf*

We previously showed that optogenetic stimulation of NPVF neurons is sufficient to promote sleep in zebrafish (Lee et al., 2017). However, it is unknown whether this phenotype is due to release of the NPVF neuropeptide or other factors within these cells, such as the fast neurotransmitter glutamate (Lee et al., 2017). To directly test the hypothesis that stimulation of *npvf*-expressing neurons promotes sleep due to release of NPVF, we optogenetically stimulated these neurons in *npvf* mutant animals. Taking advantage of the transparency of zebrafish larvae, we used a previously described non-invasive, large-scale assay that allows optogenetic stimulation of specific neuronal populations while monitoring 96 freely-behaving animals (**Figure 4 – figure supplement 1A,B**) (Singh et al., 2015). We first recorded baseline behavior for 30 minutes in the dark, and then exposed the animals to blue light for 30 minutes. Similar to previous results using animals that are homozygous wild-type for *npvf* (Lee et al., 2017), optogenetic stimulation of NPVF neurons in *Tg(npvf:ReaChR-mCitrine)*; *npvf* +/− animals resulted in reduced locomotor activity and increased sleep compared to non-transgenic *npvf* +/− sibling controls (**Figure 4 – figure supplement 1C**). In contrast, there was no significant difference between the behavior of *Tg(npvf:ReaChR-mCitrine)*; *npvf* −/− animals and their non-transgenic *npvf* −/− siblings (**Figure 4 – figure supplement 1D**). This result indicates that sleep induced by stimulation of NPVF neurons requires NPVF, and thus acts via NPVF neuropeptide/GPCR signaling.

**Figure 4.**
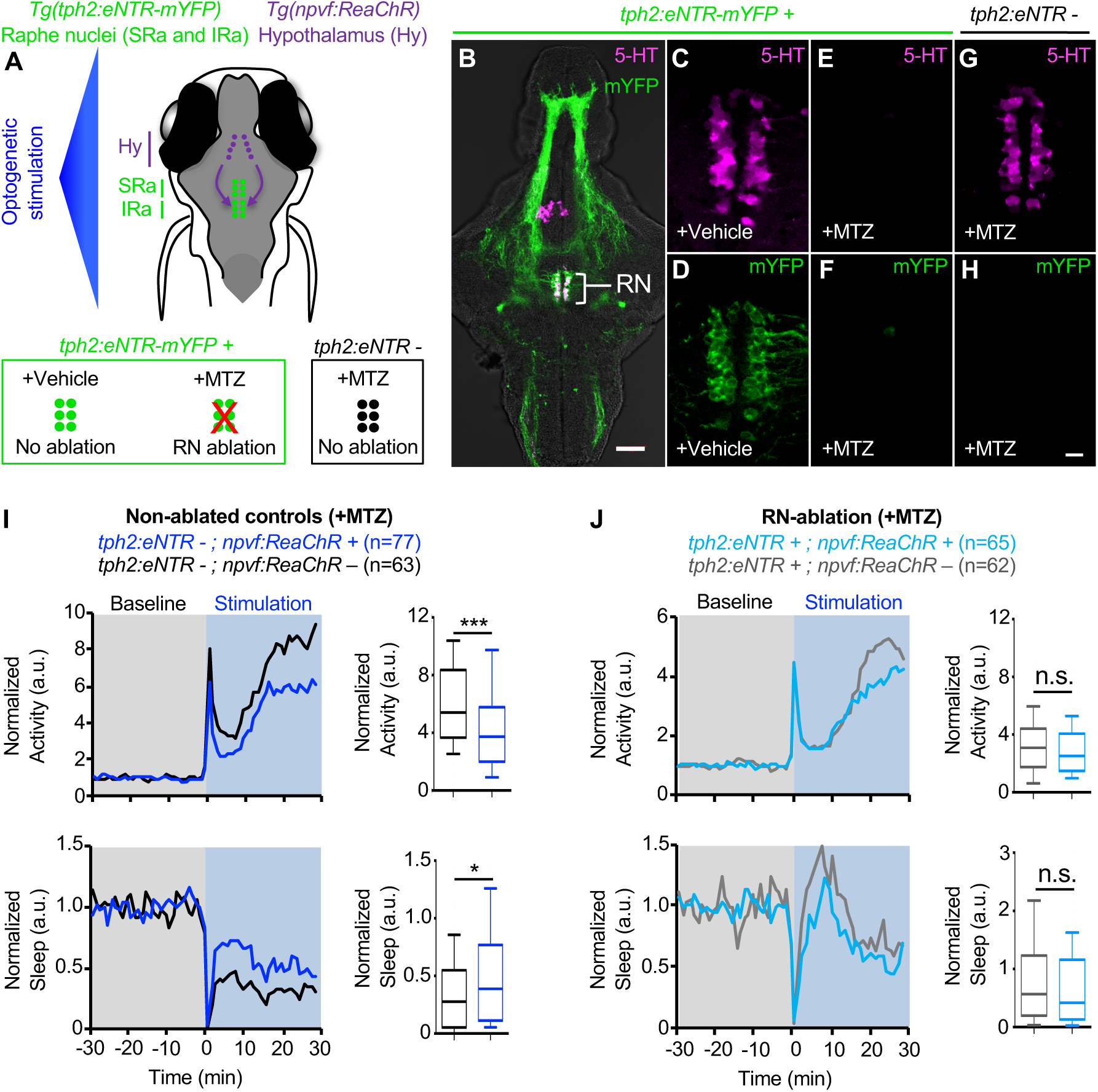
Chemogenetic ablation of the RN abolishes NPVF neuron-induced sleep. (**A**) Schematic of experiment. Treatment of animals with the small molecule MTZ results in ablation of serotonergic RN neurons in *Tg(tph2:eNTR-mYFP)* animals but not in their non-transgenic siblings. Behavior is then monitored before and during blue light exposure that stimulates *npvf*-expressing neurons in *Tg(npvf:ReaChR)* animals but not in their non-transgenic siblings. (**B**) 5-dpf *Tg(tph2:eNTR-mYFP)* zebrafish brain showing serotonergic RN neurons and some of their projections (green) and labeled with a 5-HT specific antibody (magenta). The bracketed region is magnified in (**C,D**) to show co-labeling of RN neurons with mYFP and 5-HT immunoreactivity. Treatment of *Tg(tph2:eNTR-mYFP)* animals with MTZ results in the loss of both 5-HT immunoreactivity (**E**) and YFP (**F**) in the RN, but treatment with DMSO vehicle control does not (**C,D**). MTZ treatment in *Tg(tph2:eNTR-mYFP)* negative siblings does not result in loss of RN neurons. Images are single 4 μm (**B**) and 0.6 μm (**C-H**) thick optical sections. Scale: 50 μm (**B**), 10 μm (**C-H**). (**I,J**) Normalized locomotor activity (top) and sleep (bottom) of 5-dpf *Tg(npvf:ReaChR)* (dark blue and light blue) and non-transgenic sibling controls (black and gray) zebrafish before (Baseline) and during blue light exposure (Stimulation) in *Tg(tph2:eNTR)* negative animals (**I**) and in *Tg(tph2:eNTR)* positive siblings (**J**). Box plots quantify locomotor activity and sleep during optogenetic stimulation normalized to baseline of the same genotype. n=number of animals. ns p>0.05, *p<0.05, ***p<0.005, Mann-Whitney test.

### Sleep induced by optogenetic stimulation of *npvf*-expressing neurons requires serotonergic RN neurons

We next asked whether there is a functional interaction between *npvf*-expressing neurons and the RN in promoting sleep by testing the hypothesis that NPVF neuron-induced sleep requires serotonergic RN neurons. To do so, we optogenetically stimulated *npvf*-expressing neurons in animals where the RN was chemogenetically ablated by enhanced nitroreductase (eNTR) (Mathias et al., 2014; Tabor et al., 2014). eNTR converts the inert pro-drug metronidazole (MTZ) into a cytotoxic compound that causes cell-autonomous death (Curado et al., 2007). We previously showed using *Tg*(*tph2*:*eNTR-mYFP*) animals, which express eNTR-mYFP in serotonergic RN neurons, that ablation of these neurons results in decreased sleep (Oikonomou et al., 2019). This phenotype is similar to those of both *tph2* −/− zebrafish and to mice in which the dorsal and median serotonergic RN are ablated (Oikonomou et al., 2019). To ask whether NPVF neuron-induced sleep requires the RN, we generated *Tg(npvf:ReaChR-mCitrine); Tg*(*tph2*:*eNTR-mYFP*) animals and asked whether the sedating effect of stimulating NPVF neurons is diminished in RN ablated animals (**Figure 4A,B**). Similar to our previous report (Oikonomou et al., 2019), treatment of these animals with 5 mM MTZ during 2-4 dpf resulted in near complete loss of YFP fluorescence and 5-HT immunoreactivity in the RN (**Figure 4E,F**). In contrast, treatment of these animals with DMSO vehicle control (**Figure 4C,D**), or treatment of *Tg*(*tph2*:*eNTR-mYFP*) negative siblings with MTZ (**Figure 4G,H**), did not cause loss of the RN. We found that optogenetic stimulation of MTZ-treated *Tg(npvf:ReaChR-mCitrine)* animals resulted in increased locomotor activity and decreased sleep compared to identically treated non-transgenic siblings (**Figure 4I**), indicating that MTZ treatment does not itself block NPVF neuron-induced sleep. In contrast, there was no significant difference in the behavior of MTZ-treated *Tg(npvf:ReaChR-mCitrine); Tg*(*tph2*:*eNTR-mYFP*) animals compared to their *Tg*(*tph2*:*eNTR-mYFP*) siblings (**Figure 4J**). These results indicate that the sedating effect of stimulating *npvf*-expressing neurons requires the serotonergic RN.

### Sleep induced by stimulation of *npvf*-expressing neurons requires serotonin in RN neurons

Zebrafish RN neurons produce not only serotonin, but also other factors such as GABA (Kawashima et al., 2016) that may mediate NPVF neuron-induced sleep. To distinguish between these possibilities, we next tested the hypothesis that NPVF neuron-induced sleep requires the presence of serotonin in RN neurons. To do so, we compared the effect of optogenetic stimulation of *npvf*-expressing neurons in *tph2* −/− animals, which do not synthesize serotonin in RN neurons (Oikonomou et al., 2019), to *tph2* +/− sibling controls. Similar to previous results (Lee et al., 2017), stimulation of *npvf-*expressing neurons in *Tg(npvf:ReaChR-mCitrine); tph2* +/− animals resulted in decreased locomotor activity and increased sleep (**Figure 4 – figure supplement 1E**). In contrast, there was no significant difference between the behavior of *Tg(npvf:ReaChR-mCitrine)*; *tph2* −/− animals and their non-transgenic *tph2* −/− siblings (**Figure 4 – figure supplement 1F**), consistent with the hypothesis that NPVF neuron-induced sleep requires serotonin in RN neurons.

As additional confirmation that NPVF neuron-induced sleep requires serotonin in RN neurons, we compared the effect of chemogenetic stimulation of *npvf*-expressing neurons in *tph2* −/− animals to *tph2* +/− sibling controls (**Figure 4 – figure supplement 2)**. To do so, we expressed the rat capsaicin receptor TRPV1 in NPVF neurons using *Tg(npvf:KalTA4); Tg(UAS:TRPV1-tagRFP-T*) animals (Lee et al., 2017). We previously showed that treating these animals with 2 μM capsaicin, a TRPV1 small molecule agonist, results in activation of NPVF neurons and increased sleep at night (Lee et al., 2017). Consistent our previous observations, capsaicin-treated *Tg(npvf:KalTA4); Tg(UAS:TRPV1-tagRFP-T*); *tph2* +/− animals slept more at night than their capsaicin-treated *Tg(npvf:KalTA4); tph2* +/− siblings (**Figure 4 – figure supplement 2C,F**), indicating that chemogenetic activation of NPVF neurons results in increased sleep at night in animals that produce serotonin in the RN. However, there was no significant difference in sleep at night between capsaicin-treated *Tg(npvf:KalTA4); Tg(UAS:TRPV1-tagRFP-T*); *tph2* −/− animals and their capsaicin-treated *Tg(npvf:KalTA4)*; *tph2* −/− siblings (**Figure 4 – figure supplement 2D,F**). This result is consistent with the hypothesis that NPVF neuron-induced sleep requires serotonin in RN neurons. Since 5-HT is not present in NPVF neurons (**Figure 1B-D**), our optogenetic and chemogenetic data indicate that NPVF neuron-induced sleep requires serotonin in RN neurons.

## DISCUSSION

The serotonergic RN were first implicated in sleep-wake regulation over 50 years ago, but it has long been disputed whether they act to promote sleep or wakefulness (Ursin, 2008). We and others recently addressed this controversy in both mammals and zebrafish by providing both gain- and loss-of-function evidence using genetic, pharmacological, optogenetic and chemogenetic approaches to demonstrate that the serotonergic RN promote sleep (Oikonomou et al., 2019; Venner et al., 2019). This finding agrees with invertebrate studies which showed that 5-HT signaling promotes sleep in *Drosophila* (Qian et al., 2017; Yuan et al., 2006). However, while 5-HT plays an evolutionarily conserved role in promoting sleep, the neuronal mechanism that acts upon serotonergic neurons to promote sleep was unknown. Here we show that *npvf*-expressing neurons in the dorsomedial hypothalamus, which we previously found to be sleep-promoting in zebrafish (Lee et al., 2017), densely innervate the rostral IRa, can stimulate serotonergic IRa neurons, and require 5-HT in RN neurons in order to induce sleep. These results describe a simple hypothalamic-hindbrain sleep-promoting neuronal circuit arising from the dorsomedial hypothalamus, a region previously linked to circadian regulation of wakefulness (Chou et al., 2003; Gooley et al., 2006; Mieda et al., 2006), but not to sleep. This finding suggests that re-examination of hypothalamic populations using the modern tools of neuroscience may reveal additional sleep- and wake-promoting populations.

We observed that stimulation of NPVF neurons results in increased GCaMP6s fluorescence in most serotonergic IRa neurons, although a significant minority of IRa neurons show reduced GCaMP6s fluorescence. These observations suggest that there are at least two functionally distinct neuronal populations among *tph2*-expressing IRa neurons, consistent with studies of the mammalian RN that identified molecularly distinct sub-populations with distinct efferent projection patterns and functions (Huang et al., 2019; Ren et al., 2018; Ren et al., 2019). The NPVF neuron-induced inhibition of a sub-population of IRa neurons that we observed is consistent with a previous study which reported that optogenetic stimulation of NPVF neurons resulted in decreased GCaMP6s fluorescence in IRa neurons (Madelaine et al., 2017), although the effect was small and slow compared to our data. Further studies are needed to explore the molecular and functional diversity of RN neurons in zebrafish.

The hypothalamic-hindbrain neuronal circuit that we have described can be integrated into a larger sleep-promoting network. We recently reported that epidermal growth factor receptor (EGFR) signaling is necessary and sufficient for normal sleep amounts in zebrafish, and that it promotes sleep, in part, via the NPVF system (Lee et al., 2019). We found that it does so by both promoting the expression of *npvf* and by stimulating *npvf*-expressing neurons. The EGFR ligands *egf* and *transforming growth factor alpha* are expressed in glial cells in the dorsal diencephalon, and *egfra*, the EGFR paralog that is primarily responsible for the role of EGFR signaling in sleep, is expressed in juxta-ventricular glial cells found along the hypothalamus, hindbrain, tectum and cerebellum. Taken together with the current study, these results describe a genetic and neuronal circuit spanning EGFR signaling components in glial cells, *npvf-*expressing neurons in the hypothalamus, and serotonergic RN neurons in the hindbrain.

If the EGFR-NPVF-RN sleep-promoting circuit plays a central and important role in regulating sleep, one might expect it to be evolutionarily conserved. Indeed, similar to zebrafish, EGFR signaling promotes sleep in *C. elegans* and *Drosophila* (Donlea et al., 2009; Foltenyi et al., 2007; Van Buskirk and Sternberg, 2007), and genetic experiments suggest that it does so in part via RFamide neuropeptides that may be invertebrate homologs of *npvf* (He et al., 2013; Iannacone et al., 2017; Lenz et al., 2015; Nagy et al., 2014; Nath et al., 2016; Nelson et al., 2014; Shang et al., 2013; Turek et al., 2016). Serotonin has also been shown to promote sleep in *Drosophila* (Qian et al., 2017; Yuan et al., 2006), and by analogy to our results, we hypothesize that RFamide neuropeptides such as FMRFamide (Lenz et al., 2015) may act upstream of 5-HT to promote *Drosophila* sleep. The role of EGFR signaling in mammalian sleep is less clear. Intracerebroventricular injection of epidermal growth factor (EGF) in rabbits was sufficient to increase sleep (Kushikata et al., 1998), and mice containing linked mutations in *Egfr* and *Wnt3a* (*Wingless integration site 3a*) showed abnormal circadian timing of sleep (Kramer et al., 2001). Furthermore, pharmacological inhibition or mutation of extracellular regulated kinase (ERK), which mediates EGFR signaling, was shown to result in reduced sleep in mice (Mikhail et al., 2017). The spatial expression of NPVF neurons within the hypothalamus is highly conserved between humans, rodents, and zebrafish (Lee et al., 2017; Liu et al., 2001; Ubuka et al., 2009; Yelin-Bekerman et al., 2015), as are the expression of EGFR and its ligands in zebrafish and rodent brains (Lee et al., 2019; Ma et al., 1994; Ma et al., 1992). However, NPVF has not been studied in the context of mammalian sleep, so further studies are required to determine whether the EGFR/NPVF/RN circuit described in zebrafish is conserved in mammals.

## MATERIALS AND METHODS

### KEY RESOURCE TABLE

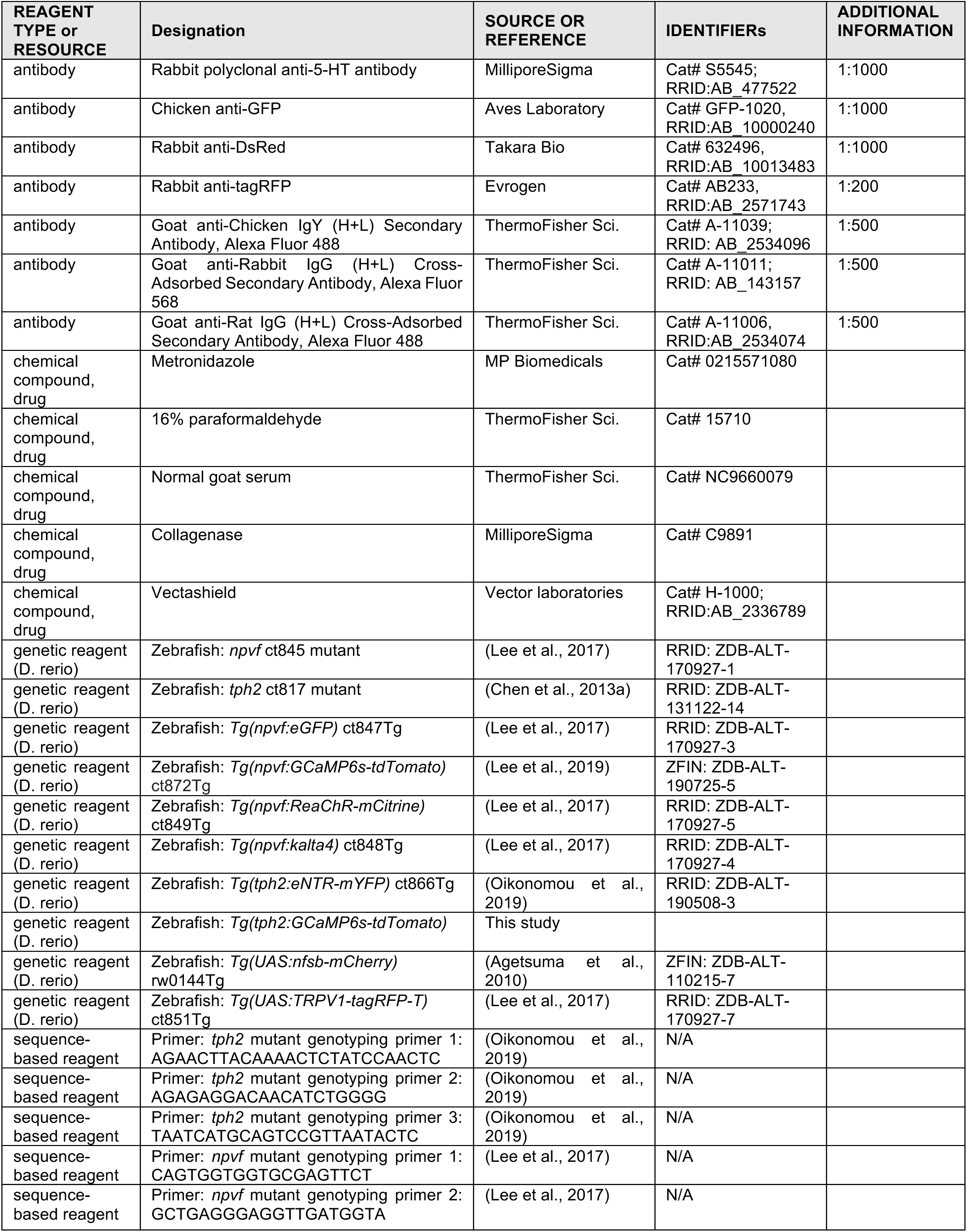
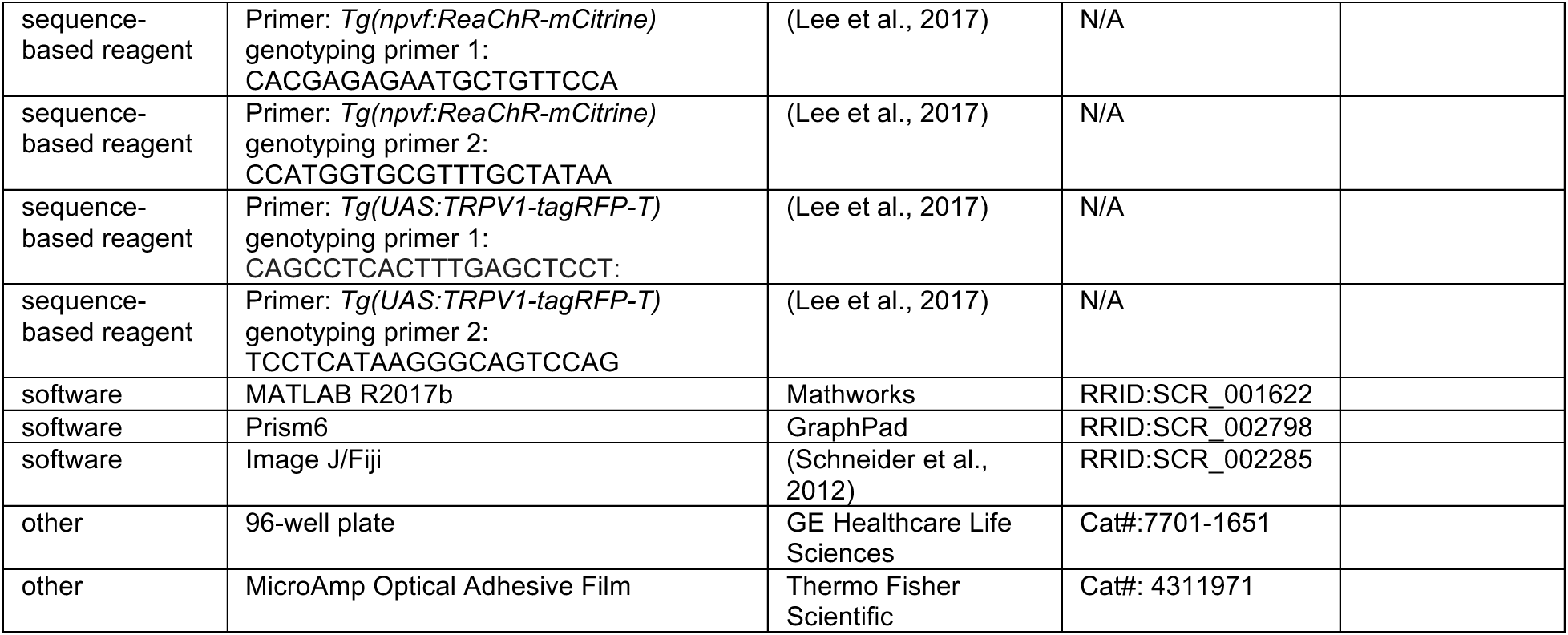

### EXPERIMENTAL MODEL AND SUBJECT DETAILS

Animal husbandry and all experimental procedures involving zebrafish were performed in accordance with the California Institute of Technology Institutional Animal Care and Use Committee (IACUC) guidelines and by the Office of Laboratory Animal Resources at the California Institute of Technology (animal protocol 1580). All experiments used zebrafish between 5 and 6 dpf. Sex is not yet defined at this stage of development. Larvae were housed in petri dishes with 50 animals per dish. E3 medium (5 mM NaCl, 0.17 mM KCl, 0.33 mM CaCl_2_, 0.33 mM MgSO_4_) was used for housing and experiments. All lines were derived from the TLAB hybrid strain. Unless otherwise indicated, for experiments using mutant animals, heterozygous and homozygous mutant adult animals were mated, and their homozygous mutant and heterozygous mutant progeny were compared to each other to minimize variation due to genetic background. For experiments using transgenic animals, heterozygous transgenic animals were outcrossed to non-transgenic animals of the parental TLAB strain, and transgenic heterozygote progeny were compared to their non-transgenic siblings. Behavioral experiments were performed unbiased to genotype, with animals genotyped by PCR after each experiment was complete.

#### Transgenic and mutant animals

The *Tg(npvf:eGFP)* ct847Tg (Lee et al., 2017), *Tg(npvf:ReaChR-mCitrine)* ct849Tg (Lee et al., 2017), *Tg(npvf:kalta4)* ct848Tg (Lee et al., 2017), *Tg(tph2:eNTR-mYFP)* ct866Tg (Oikonomou et al., 2019), *Tg(UAS:nfsb-mCherry)* rw0144Tg (Agetsuma et al., 2010), *Tg(UAS:TRPV1-tagRFP-T)* ct851Tg (Lee et al., 2017), *npvf* ct845 mutant (Lee et al., 2017), and *tph2* ct817 mutant (Chen et al., 2013a) lines have been previously described.

To generate the *Tg(tph2:GCaMP6s-P2A-NLS-tdTomato)* line we cloned the *tph2* promoter (Oikonomou et al., 2019) upstream of GCaMP6s (Chen et al., 2013b) followed by a P2A sequence, which generates a self-cleaving peptide (Kim et al., 2011), and NLS-tdTomato. Stable transgenic lines were generated using the Tol2 method (Urasaki et al., 2006).

#### Immunohistochemistry

Samples were fixed in 4% PFA/4% sucrose in PBS overnight at 4°C and then washed with 0.25% Triton X-100/PBS (PBTx). Immunolabeling was performed using dissected brains because this allows for superior antibody penetration. Dissected brains were incubated for 1 hour in 1 mg/mL collagenase (C9891, MilliporeSigma, St. Louis, Missouri, USA) and blocked overnight in 2% NGS/2% DMSO in PBTx at 4°C. Incubation with rabbit anti-5-HT (1:1000; S5545, MilliporeSigma, Burlington, MA, USA), chicken anti-GFP (1:1000, GFP-1020, Aves Laboratory, Davis, CA, USA), and rabbit anti-DsRed (1:1000, Takara Bio, Mountainview, CA, USA) primary antibodies was performed in blocking solution overnight at 4°C. Incubation with goat anti-rabbit IgG Alexa Fluor 568, goat anti-chicken IgY Alexa Fluor 488, and goat anti-rat IgG Alexa Fluor 488 (all 1:500, ThermoFisher Sci., Waltham, MA, USA) secondary antibodies was performed in blocking solution overnight at 4°C. Samples were mounted in Vectashield (H-1000; Vector Laboratories, Burlingame, CA, USA) and imaged using a Zeiss LSM 880 confocal microscope (Zeiss, Oberkochen, Germany).

#### Two-photon optogenetic stimulation and GCaMP6s imaging

At 6 dpf, animals were paralyzed by immersion in 1 mg/ml α-bungarotoxin (2133, Tocris, Bristol, UK) dissolved in E3, embedded in 1.5% low melting agarose (EC-202, National Diagnostics, Atlanta, GA, United States) and imaged using a 20x water immersion objective on a Zeiss LSM 880 microscope equipped with a two-photon laser (Chameleon Coherent, Wilsonville, OR, USA). Laser power coming out of the objective was quantified using a power meter (PM121D, ThorLabs, Newton, NJ, USA). For GCaMP6s imaging, a region of interest (ROI) that encompassed the *npvf-* or *tph2*-expressing neuronal somas was defined based on nuclear localized tdTomato, which was stoichiometrically co-expressed with GCaMP6s. GCaMP6s fluorescence intensity was quantified using Image J (Schneider et al., 2012). **Figure 2 – figure supplement 1F** provides an example of how ROI were defined. GCaMP6s fluorescence was excited using a 920 nm two-photon laser (Chameleon Coherent, Wilsonville, OR, USA) at 8 mW, imaged in a 512 × 256 pixel frame (1.27 s per frame, pixel size=0.55 μm, pixel dwell time=2.07 μs) for 150 frames (190.5 s). For optogenetic stimulation of NPVF neurons, a 150 × 100 pixel region that encompassed the NPVF neuronal somas was illuminated using the 920 nm two-photon laser at 38 mW. Ten pulses were applied over 3.72 s using the bleaching function at 2.7 Hz per pixel. The time between the final stimulation pulse and initiation of post-stimulation imaging was 0.6 s, and was due to the computer registering coordinate information with the scan device (∼0.4 s) and for the non-descanned detector to turn off (∼0.2 s). GCaMP6s fluorescence was then imaged again for another 150 frames before another stimulation trial. Three stimulation trials were performed on each fish, and the average value from these trials was calculated for each neuron. Baseline fluorescence (F_0_) for each trial was defined as the average value from 30 frames immediately preceding the stimulation, and 35 (**Figure 2D-F**) or 30 (**Figure 2 – figure supplement 1G-I**) frames post stimulation were used to measure the normalized change in fluorescence (ΔF/F = (F−F_0_)/(F_0_ −F_background_). F_background_ was defined by average fluorescence intensity during 30 frames prior to optogenetic stimulation in a GCaMP-negative background region approximately 40 μm from the nearest NPVF neuronal soma. Five *Tg(npvf:ReaChR);Tg(npvf:GCaMP6s-tdTomato)* animals, with approximately 17 NPVF neurons per animal, were analyzed for **Figure 2 – figure supplement 1**. Data from four *Tg(npvf:ReaChR);Tg(tph2:GCaMP6s-tdTomato)* and four *Tg(npvf:eGFP);Tg(tph2:GCaMP6s-tdTomato)* animals, with approximately 30 IRa neurons analyzed per animal, is shown in **Figure 2 – figure supplement 1**.

#### Sleep/wake behavioral analysis

Sleep/wake analysis was performed as previously described (Chiu et al., 2016). Larvae were raised on a 14:10 h light:dark (LD) cycle at 28.5°C with lights on at 9 a.m. and off at 11 p.m. Dim white light was used to raise larvae for optogenetic experiments to prevent stimulation of ReaChR by ambient light. Individual larvae were placed into each well of a 96-well plate (7701-1651, Whatman, Pittsburgh, PA, United States) containing 650 μl of E3 embryo medium. Locomotor activity was monitored using a videotracking system (Viewpoint Life Sciences, Lyon, France) with a Dinion one-third inch Monochrome camera (Dragonfly 2, Point Grey, Richmond, Canada) fitted with a variable-focus megapixel lens (M5018-MP, Computar, Cary, NC, United States) and infrared filter. The movement of each larva was recorded using the quantization mode. The 96-well plate and camera were housed inside a custom-modified Zebrabox (Viewpoint Life Sciences) that was continuously illuminated with infrared light. The 96-well plate was housed in a chamber filled with recirculating water to maintain a constant temperature of 28.5°C. The parameters used for movement detection were: detection threshold, 15; burst, 29; freeze, 3; bin size, 60 s, which were determined empirically. Data were analyzed using custom Perl and Matlab (Mathworks, Natick, MA, United States) scripts, which conform to the open source definition.

#### Optogenetic stimulation of freely behaving animals

Optogenetic behavioral experiments were performed as described (Singh et al., 2015). These experiments used a videotracking system with a custom array containing three sets of blue LEDs (470 nm, MR-B0040-10S, Luxeon V-star, Brantford, Canada) mounted 15 cm above and 7 cm away from the center of the 96-well plate to ensure uniform illumination. The LEDs were controlled using a custom-built driver and software written in BASIC stamp editor. A power meter (1098293, Laser-check, Santa Clara, CA, USA) was used before each experiment to verify uniform light intensity (∼400 μW at the surface of the 96-well plate). In the afternoon of the fifth day of development, single larvae were placed into each well of a 96-well plate and placed in the videotracker in the dark. Larvae were exposed to blue light for 30 minutes for each of 3 trials beginning at 12:30 am, 3:00 am, 5:30 am. Behavior was monitored for 30 minutes before and after light onset. Light onset induces a startle response, which causes a short burst of locomotor activity. For this reason, we excluded five minutes of behavioral recording centered at the peak of blue light onset from analysis. Data was normalized by dividing the locomotor activity or sleep of each animal during light exposure by the average baseline locomotor activity or sleep of all animals of the same genotype. For baseline, we used a time period equal in length to blue light exposure, but prior to light onset.

#### Chemogenetic ablation

Animals were treated with 5 mM metronidazole (MTZ) (0215571080, MP Biomedicals, Santa Ana, CA, USA) diluted in E3 medium containing 0.1% DMSO, starting in the afternoon at 2 dpf, and refreshed every 24 hours. Animals were kept in dim light during the day to prevent MTZ photodegradation. On the evening at 4-dpf, the animals were rinsed 3 times in E3 medium, moved into petri dishes to allow for recovery overnight, and then transferred to 96-well plates on the morning of 5-dpf. Reported data is from the 5th night of development.

#### Chemogenetic stimulation

Neuronal activation using TRPV1 was performed as described (Lee et al., 2017) with some modifications. *Tg(npvf:KalTA4); tph2+/−, Tg(npvf:KalTA4); tph2−/−, Tg(npvf:KalTA4); Tg(UAS:TRPV1-tagRFP-T); tph2+/−*, and *Tg(npvf:KalTA4);Tg(UAS:TRPV1-tagRFP-T); tph2−/−* siblings were immersed in 2 μM capsaicin at ∼100 h post-fertilization (hpf). Capsaicin powder (M2028, Sigma, St. Louis, Missouri, USA) was dissolved in DMSO to prepare a 100 mM stock solution that was stored in aliquots at −20 °C. Capsaicin working solutions were prepared just before each experiment by diluting the stock solution in E3 medium. All treatments contained a final concentration of 0.002% DMSO. Behavioral analysis was performed from 5 dpf until 6 dpf blinded to the genotype.

### QUANTIFICATION AND STATISTICAL ANALYSIS

For all behavioral experiments the unit of analysis for statistics is a single animal. For optogenetic imaging experiments, the unit of analysis for statistics is the average of three optogenetic stimulation trials for a single neuron. The number of neurons and animals whose data are shown in a panel are either shown in the figure or stated in the figure legend. Line graphs in **Figure 3A-D** and **Figure 4 – figure supplement 2A-D** represent mean, and were generated from raw data that was smoothed over 1 hour bins in 10 minute intervals. The significance threshold was set to P<0.05, and P values were adjusted for multiple comparisons where appropriate. Normality tests (D’Agostino & Pearson omnibus normality test) found that most datasets were not normally distributed so we used non-parametric tests for statistical analyses (Mann-Whitney test for two unpaired groups). For comparison of differences between groups with two-factor designs, we used Two-Way ANOVA with Holm-Sidak correction for multiple comparisons (**Figure 3,4**). Normality tests (D’Agostino & Pearson omnibus normality test) for the data in **Figures 3** and **Figure 4 – figure supplement 2** demonstrated that half of values were normally distributed, suggesting that Two-Way ANOVA would be an appropriate statistical test; ANOVA analyses are robust to even large deviation from normality when samples sizes are appropriately large enough.

For box plots, the box extends from the 25^th^ to the 75^th^ percentile with the median marked by a horizontal line through the box. The lower and upper whiskers extend to the 10^th^ and 90^th^ percentile, respectively. Data points outside the lower and upper whiskers were not shown in the graphs to facilitate data presentation but were included in statistical analyses. Statistical analyses were performed using Prism 6 (GraphPad Software, San Diego, CA, USA).

## DATA AND CODE AVAILABILITY

Source figures are publicly available at https://elifesciences.org/articles/25727 (Lee et al., 2017). All input data used to generate most figures is available as source data associated with this manuscript.

**Figure 1-Source Data 1:** Confocal stack of a brain from a 6-dpf *Tg(npvf:eGFP)* animal immunostained against eGFP and 5-HT.

**Figure 1-Supplement1-Source Data 1:** Confocal stacks of a brain from a 6-dpf *Tg(npvf:KalTA4);Tg(uas:nfsb-mCherry);Tg(tph2:eNTR-mYFP)* animal immunostained against YFP and mCherry.

**Figure 2-Source Data 1: (A)** Confocal image of a live 6-dpf *Tg(npvf:ReaChR-mCitrine); Tg(tph2:GCaMP6s-tdTomato)* animal showing GCaMP6s and tdTomato fluorescence in IRa neurons. Green and magenta fluorescence in the eyes is due to autofluorescence. (**C**) Average GCaMP6s fluorescence in IRa neurons for 10 frames before (Pre) and after (Post) optogenetic stimulation of NPVF neurons. (**D,E**) Input data and statistical analysis for GCaMP6s fluorescence of IRa neurons plotted as a function of time, showing pre-stimulation and post-stimulation evoked responses for *Tg(tph2:GCaMP6s-tdTomato)* animals that also contain a *Tg(npvf:eGFP)* (**D,F**) or *Tg(npvf:ReaChR-mCitrine)*.

**Figure 2-Supplement1-Source Data 1: (A)** Confocal image of a live 6-dpf *Tg(npvf:ReaChR); Tg(npvf:GCaMP6s-tdTomato)* animal showing GCaMP6s and tdTomato fluorescence in NPVF neurons. Green and magenta fluorescence in the eyes is due to autofluorescence. (**H**) GCaMP6s and mCitrine **Δ**F/F values for individual NPVF neurons (86 neurons, 5 animals) are plotted as function of time, showing pre-stimulation and post-stimulation evoked responses. (**I**) Statistical analysis for GCaMP6s and mCitrine **Δ**F/F values one frame after stimulation for *Tg(npvf:GCaMP6s-tdTomato)* animals that also contain a *Tg(npvf:ReaChR)* (blue) or *Tg(npvf:eGFP)* (black) transgene. n=number of neurons.

**Figure 3-Source Data 1:** Input data and statistical analysis for the average sleep for *npvf* +/−; *tph2* +/−, *npvf* −/−; *tph2* +/−, *npvf* +/−; *tph2* −/−, and *npvf* −/−; *tph2* −/− siblings.

**Figure 4-Source Data 1: (B)** Confocal image of an immunostained 5-dpf *Tg(tph2:eNTR-mYFP)* zebrafish brain showing serotonergic RN neurons and some of their projections and labeled with a 5-HT specific antibody. (**I-J**) Input data and statistical analysis for normalized locomotor activity and sleep of 5-dpf *Tg(npvf:ReaChR)* and non-transgenic sibling controls (black and gray) zebrafish before (Baseline) and during blue light exposure (Stimulation) in *Tg(tph2:eNTR)* negative animals (**I**) and in *Tg(tph2:eNTR)* positive siblings (**J**). Box plots quantify locomotor activity and sleep during optogenetic stimulation normalized to baseline of the same genotype. n=number of animals.

**Figure 4-Supplement1-Source Data 1: (A)** Confocal image of a live 5-dpf *Tg(npvf:ReaChR-mCitrine)* animal. NPVF neurons in the hypothalamus are labeled with the *npvf:ReaChR-mCitrine* transgene. Green fluorescence in the eyes is due to autofluorescence. (**C,D**) Input data and statistical analysis for normalized locomotor activity and sleep of *Tg(npvf:ReaChR)* and non-transgenic sibling control animals before (Baseline) and during exposure to blue light (Stimulation) in *npvf* +/− (**C**) and *npvf* −/− (**D**) animals. (**E,F**) Same as (**C,D**) but with *tph2* mutant rather than *npvf* mutant. Because the animals see the blue light, they exhibit a brief startle at light onset that is excluded from analysis, followed by a gradual increase in activity that plateaus after ∼15 minutes. Box plots quantify locomotor activity and sleep for each animal during optogenetic stimulation normalized to the baseline of all animals the same genotype. Stimulation of *npvf*-expressing neurons decreases locomotor activity and increases sleep compared to non-transgenic sibling controls in *npvf* +/− animals (**C**) but not in *npvf* −/− siblings (**D**), and in *tph2* +/− animals (**E**) but not in *tph2* −/− siblings (**F**). n=number of animals.

**Figure 4-Supplement2-Source Data 2: (A)** Input data and statistical analysis for sleep associated with 5-6dpf *Tg(npvf:KalTA4); tph2+/−, Tg(npvf:KalTA4);tph2-*/-, *Tg(npvf:KalTA4);Tg(UAS:TRPV1-TagRFP-T);tph2*+/−, and *Tg(npvf:KalTA4);Tg(UAS:TRPV1-TagRFP-T);tph2−/−* (light blue) siblings treated with 2 μM capsaicin.

## ACKNOWLEDGEMENTS

We thank members of the Prober lab for helpful discussions; Uyen Pham, Chris Cook, and Hannah Hurley for zebrafish husbandry assistance; and Andres Collazo, Giada Spigolon, Andrey Andreev and the Beckman Institute Biological Imaging Facility for 2-photon imaging assistance. This work was supported by grants from the NIH (DAL: K99NS097683, F32NS084769; GO: F32NS084769; DAP: NS070911, NS101158), a NARSAD Young Investigator Grant (DAL: 25392) and a Caltech BBE Postdoctoral Fellowship to DAL. The authors declare no competing interests.

## AUTHOR CONTRIBUTIONS

DAL and DAP designed experiments. DAL, TC and YH performed experiments and analysis. GO generated *tph2* transgenic animals and Matlab code. DAP supervised the project. DAL and DAP wrote the paper with input from GO.

## AUTHOR ORCIDs

Daniel A Lee, http://orcid.org/0000-0001-7411-2740

Grigorios Oikonomou, http://orcid.org/0000-0001-6797-7375

David A Prober, http://orcid.org/0000-0002-7371-4675

## ETHICS

Animal experimentation: This study was performed in strict accordance with the recommendations in the Guide for the Care and Use of Laboratory Animals of the National Institutes of Health. All experiments were performed using standard protocols (Westerfield, 1993) in accordance with the California Institute of Technology Institutional Animal Care and Use Committee guidelines.

---

**Figure 1 – figure supplement 1.**
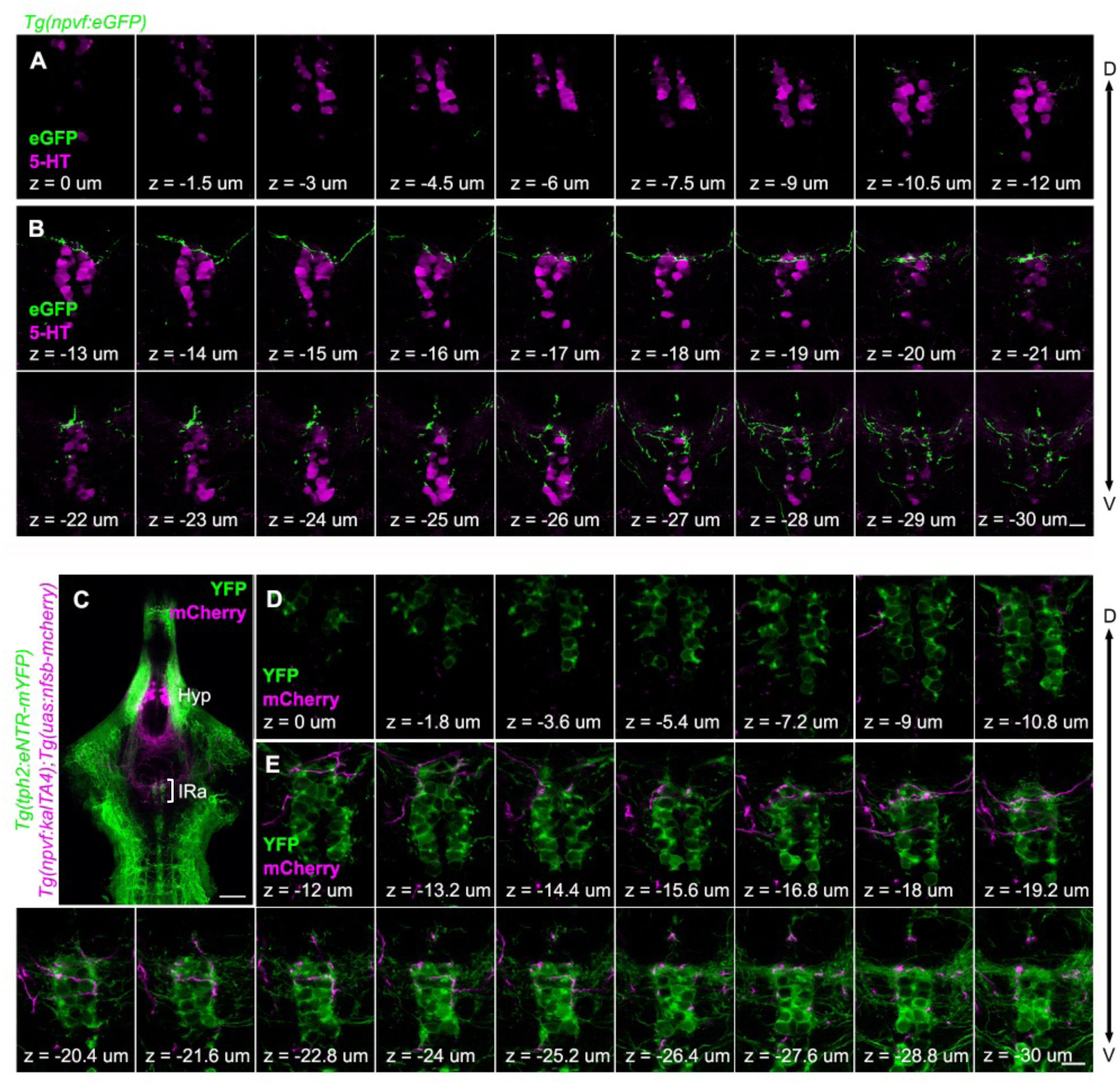
Projections of hypothalamic NPVF neurons to the serotonergic IRa shown in single optical sections. Related to Figure 1. (**A,B**) Serial optical sections (0.5 μm thick) in the hindbrain of a 6-dpf *Tg(npvf:eGFP)* animal labeled with a 5-HT-specific antibody (magenta), which were used to generate the image shown in Figure 1K. Fibers from *npvf*-expressing neurons (green) do not innervate SRa soma (**A**) but do innervate IRa soma (**B**). (**C**) A 4 μm thick optical section of a brain from a 6-dpf *Tg(npvf:KalTA4); Tg(uas:nfsb-mCherry); Tg(tph2:eNTR-mYFP)* animal. White bracket indicates the IRa and is magnified in panels (**D,E**), which show 0.6 μm thick serial optical sections. Fibers from *npvf*-expressing neurons (magenta) do not innervate SRa neurons (green, **D**) but do innervate IRa neurons and their fibers (green, **E**). Hyp, hypothalamus, IRa, inferior raphe; D, dorsal; V, ventral. Scale: 50 μm (**C**), 10 μm (**B,E**).

**Figure 2 – figure supplement 1.**
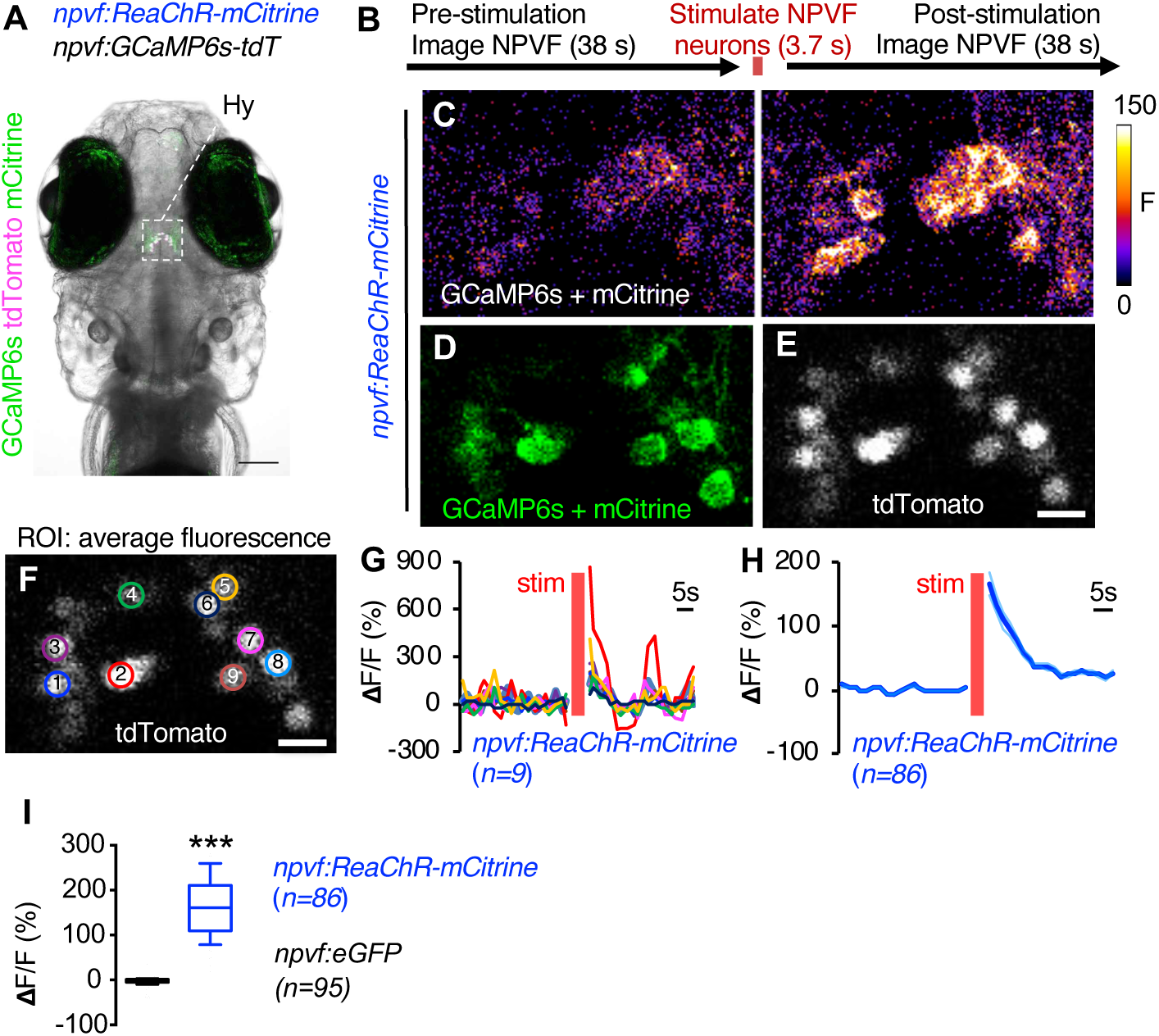
Validation of two-photon-induced optogenetic stimulation of NPVF neurons. Related to Figure 2. 6-dpf *Tg(npvf:ReaChR); Tg(npvf:GCaMP6s-tdTomato)* animals are analyzed for GCaMP6s fluorescence levels in NPVF neurons before and after optogenetic stimulation of NPVF neurons. (**A**) Example of a 6-dpf *Tg(npvf:ReaChR); Tg(npvf:GCaMP6s-tdTomato)* animal showing GCaMP6s and tdTomato fluorescence in NPVF neurons. Green and magenta signals in the eyes are due to autofluorescence. White box indicates the region of the hypothalamus that was stimulated and analyzed in subsequent panels. (**B**) Baseline GCaMP6s fluorescence in NPVF neurons was first recorded using two-photon 920 nm light at 8 mW laser power for 38 s (30 frames). NPVF neurons were then optogentically stimulated using 920 nm light at 38 mW at 2.7 Hz for 3.7 s. GCaMP6s fluorescence in the IRa was then immediately imaged again using 8 mW laser power for 38 s (30 frames). (**C**) GCaMP6s fluorescence in NPVF neurons recorded one frame before (Pre) and after (Post) optogenetic stimulation of NPVF neurons. (**D,E**) Structural images showing the location of each NPVF neuron that were obtained by averaging 10 frames in the green (**D**) or red (**E**) channel. (**F**) Regions of interest (ROI) to quantify GCaMP6s fluorescence in individual NPVF neurons were defined based on nuclear tdTomato fluorescence. (**G,H**) GCaMP6s and mCitrine **Δ**F/F values for individual NPVF neurons (**G**, 9 neurons, 1 animal) and mean ± SEM GCaMP6s and mCitrine **Δ**F/F values (**H**, 86 neurons, 5 animals) are plotted as function of time, showing pre-stimulation and post-stimulation evoked responses. (**I**) Box plots quantify GCaMP6s and mCitrine **Δ**F/F values one frame after stimulation for *Tg(npvf:GCaMP6s-tdTomato)* animals that also contain a *Tg(npvf:ReaChR)* (blue) or *Tg(npvf:eGFP)* (black) transgene. n=number of neurons. ***p<0.0005, Mann-Whitney test. Scale: 100 μm (**A**), 10 μm (**C-F**).

**Figure 4 – figure supplement 1.**
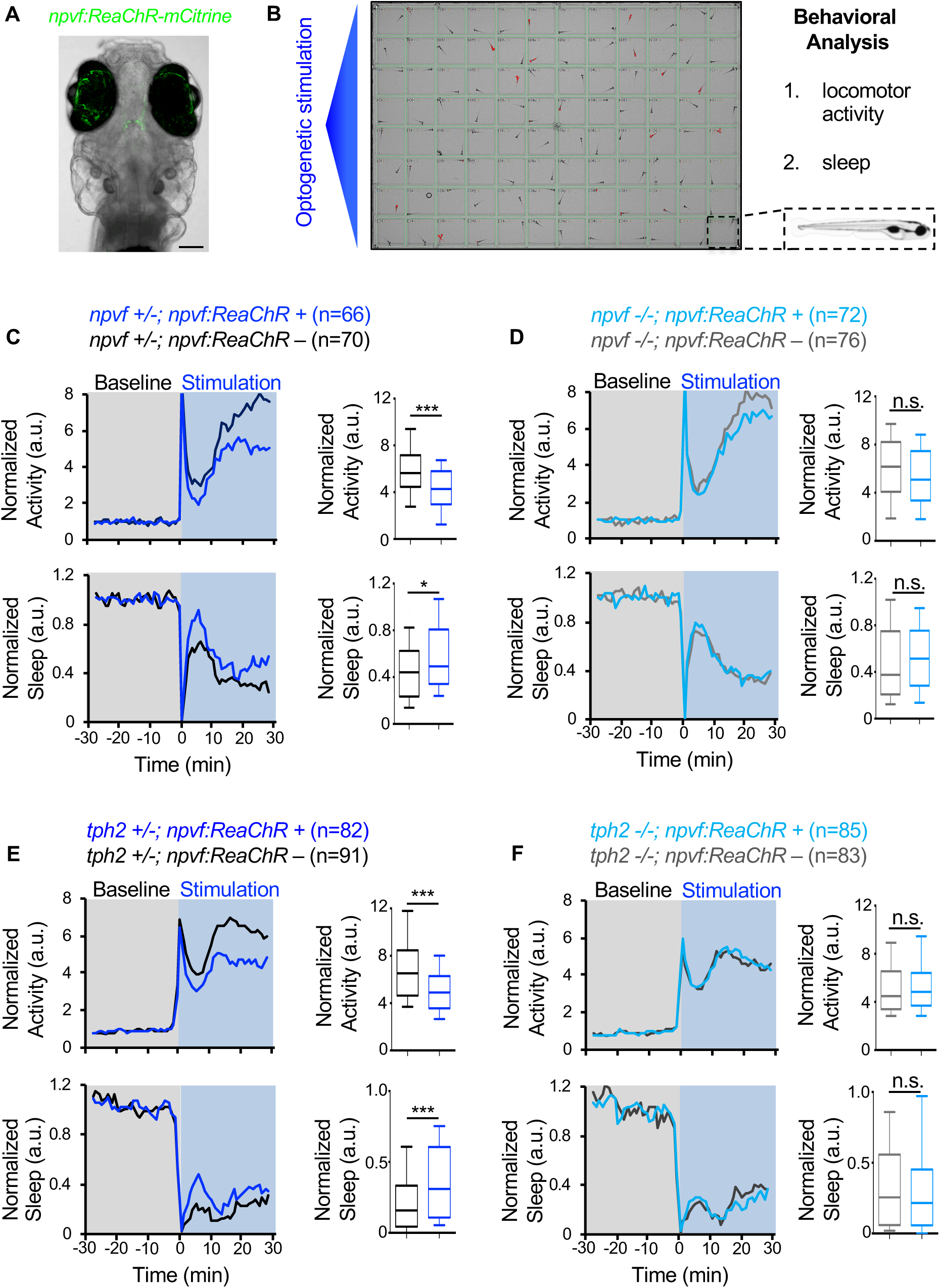
Sleep induced by optogenetic stimulation of *npvf*-expressing neurons is abolished in *npvf* and *tph2* mutant animals. Related to Figure 4. (**A,B**) 5-dpf *Tg(npvf:ReaChR-mCitrine)* animals and their non-transgenic siblings were analyzed for changes in locomotor activity and sleep in a 96-well plate optogenetic assay. NPVF neurons in the hypothalamus are labeled with the *npvf:ReaChR-mCitrine* transgene. Green fluorescence in the eyes is due to autofluorescence. (**C,D**) Normalized locomotor activity (top) and sleep (bottom) of *Tg(npvf:ReaChR)* (dark blue and light blue) and non-transgenic sibling control (black and gray) animals before (Baseline) and during exposure to blue light (Stimulation) in *npvf* +/− (**C**) and *npvf* −/− (**D**) animals. (**E,F**) Same as (**C,D**) but with *tph2* mutant rather than *npvf* mutant. Because the animals see the blue light, they exhibit a brief startle at light onset that is excluded from analysis, followed by a gradual increase in activity that plateaus after ∼15 minutes. Box plots quantify locomotor activity and sleep for each animal during optogenetic stimulation normalized to the baseline of all animals the same genotype. Stimulation of *npvf*-expressing neurons decreases locomotor activity and increases sleep compared to non-transgenic sibling controls in *npvf* +/− animals (**C**) but not in *npvf* −/− siblings (**D**), and in *tph2* +/− animals (**E**) but not in *tph2* −/− siblings (**F**). n=number of animals. ns p>0.05, *p<0.05, ***p<0.005, Mann-Whitney test.

**Figure 4 – figure supplement 2.**
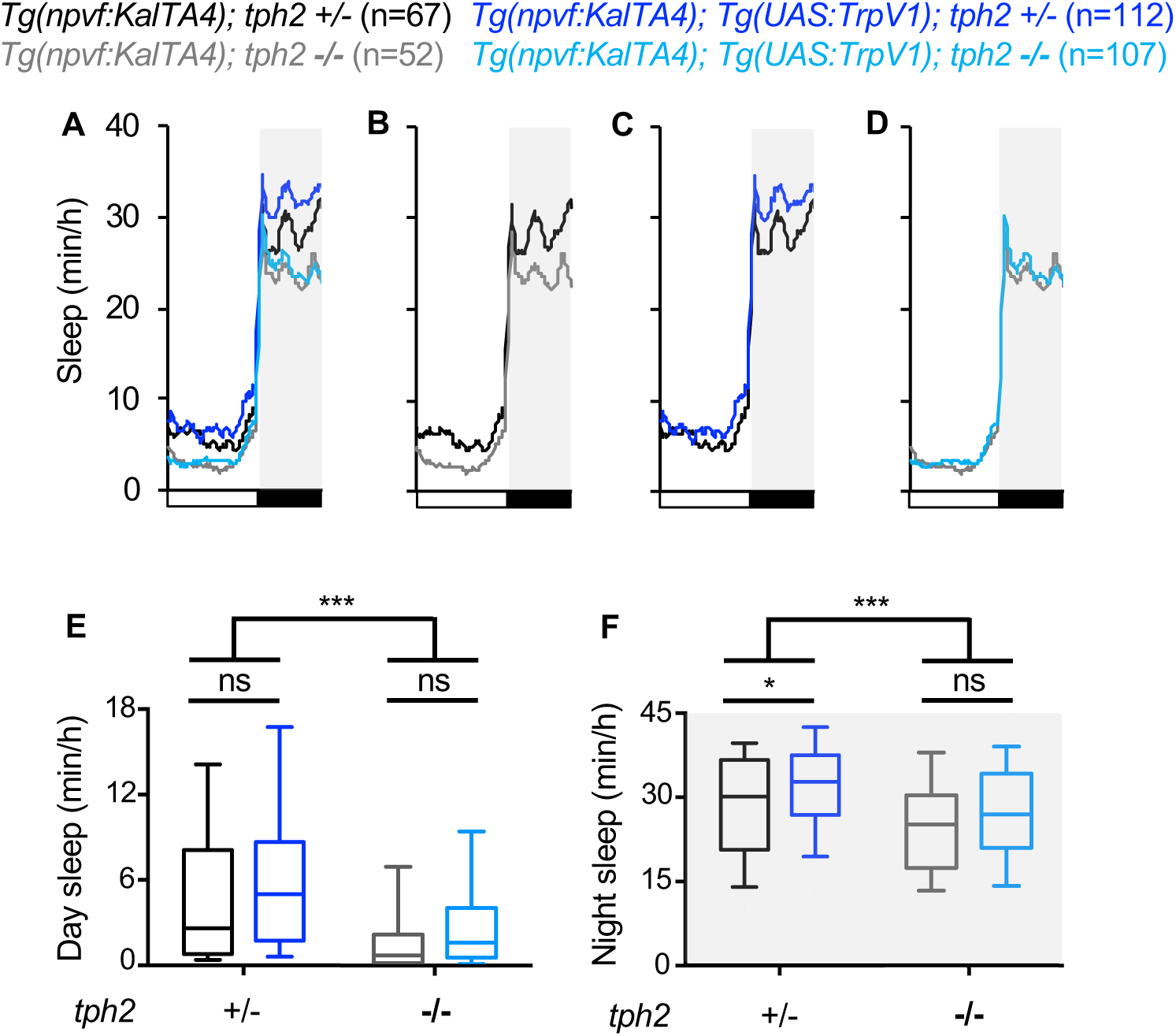
Sleep induced by chemogenetic stimulation of *npvf*-expressing neurons is abolished in *tph2* mutant animals. Related to Figure 4. (A) Average sleep for 5-dpf *Tg(npvf:KalTA4); tph2+/−* (black)*, Tg(npvf:KalTA4);tph2-*/- (gray), *Tg(npvf:KalTA4);Tg(UAS:TRPV1-TagRFP-T);tph2*+/− (dark blue), and *Tg(npvf:KalTA4);Tg(UAS:TRPV1-TagRFP-T);tph2−/−* (light blue) siblings treated with 2 μM capsaicin. Box plots quantify sleep during the day (**E**) and night (**F**). White boxes indicate day. Black boxes and gray shading indicate night. n=number of animals. ns p>0.05, *p<0.05,***p<0.005, Two-way ANOVA with Holm-Sidak test.

## REFERENCES

Agetsuma, M., Aizawa, H., Aoki, T., Nakayama, R., Takahoko, M., Goto, M., Sassa, T., Amo, R., Shiraki, T., Kawakami, K., et al. (2010). The habenula is crucial for experience-dependent modification of fear responses in zebrafish. Nat Neurosci 13, 1354–1356.

Bringmann, H. (2018). Sleep-Active Neurons: Conserved Motors of Sleep. Genetics 208, 1279–1289.

Chen, S., Oikonomou, G., Chiu, C.N., Niles, B.J., Liu, J., Lee, D.A., Antoshechkin, I., and Prober, D.A. (2013a). A large-scale in vivo analysis reveals that TALENs are significantly more mutagenic than ZFNs generated using context-dependent assembly. Nucleic acids research 41, 2769–2778.

Chen, T.W., Wardill, T.J., Sun, Y., Pulver, S.R., Renninger, S.L., Baohan, A., Schreiter, E.R., Kerr, R.A., Orger, M.B., Jayaraman, V., et al. (2013b). Ultrasensitive fluorescent proteins for imaging neuronal activity. Nature 499, 295–300.

Cheng, R.K., Krishnan, S., and Jesuthasan, S. (2016). Activation and inhibition of tph2 serotonergic neurons operate in tandem to influence larval zebrafish preference for light over darkness. Sci Rep 6, 20788.

Chiu, C.N., Rihel, J., Lee, D.A., Singh, C., Mosser, E.A., Chen, S., Sapin, V., Pham, U., Engle, J., Niles, B.J., et al. (2016). A Zebrafish Genetic Screen Identifies Neuromedin U as a Regulator of Sleep/Wake States. Neuron 89, 842–856.

Chou, T.C., Scammell, T.E., Gooley, J.J., Gaus, S.E., Saper, C.B., and Lu, J. (2003). Critical role of dorsomedial hypothalamic nucleus in a wide range of behavioral circadian rhythms. J Neurosci 23, 10691–10702.

Curado, S., Anderson, R.M., Jungblut, B., Mumm, J., Schroeter, E., and Stainier, D.Y. (2007). Conditional targeted cell ablation in zebrafish: a new tool for regeneration studies. Dev Dyn 236, 1025–1035.

Dahlstroem, A., and Fuxe, K. (1964). Evidence for the Existence of Monoamine-Containing Neurons in the Central Nervous System. I. Demonstration of Monoamines in the Cell Bodies of Brain Stem Neurons. Acta Physiol Scand Suppl, SUPPL 232:231–255.

Donlea, J.M., Ramanan, N., and Shaw, P.J. (2009). Use-dependent plasticity in clock neurons regulates sleep need in Drosophila. Science 324, 105–108.

Foltenyi, K., Greenspan, R.J., and Newport, J.W. (2007). Activation of EGFR and ERK by rhomboid signaling regulates the consolidation and maintenance of sleep in Drosophila. Nat Neurosci 10, 1160–1167.

Gooley, J.J., Schomer, A., and Saper, C.B. (2006). The dorsomedial hypothalamic nucleus is critical for the expression of food-entrainable circadian rhythms. Nat Neurosci 9, 398–407.

He, C., Yang, Y., Zhang, M., Price, J.L., and Zhao, Z. (2013). Regulation of sleep by neuropeptide Y-like system in Drosophila melanogaster. PLoS One 8, e74237.

Hinuma, S., Shintani, Y., Fukusumi, S., Iijima, N., Matsumoto, Y., Hosoya, M., Fujii, R., Watanabe, T., Kikuchi, K., Terao, Y., et al. (2000). New neuropeptides containing carboxy-terminal RFamide and their receptor in mammals. Nat Cell Biol 2, 703–708.

Huang, K.W., Ochandarena, N.E., Philson, A.C., Hyun, M., Birnbaum, J.E., Cicconet, M., and Sabatini, B.L. (2019). Molecular and anatomical organization of the dorsal raphe nucleus. Elife 8.

Iannacone, M.J., Beets, I., Lopes, L.E., Churgin, M.A., Fang-Yen, C., Nelson, M.D., Schoofs, L., and Raizen, D.M. (2017). The RFamide receptor DMSR-1 regulates stress-induced sleep in C. elegans. Elife 6.

Kawashima, T., Zwart, M.F., Yang, C.T., Mensh, B.D., and Ahrens, M.B. (2016). The Serotonergic System Tracks the Outcomes of Actions to Mediate Short-Term Motor Learning. Cell 167, 933–946 e920.

Kim, J.H., Lee, S.R., Li, L.H., Park, H.J., Park, J.H., Lee, K.Y., Kim, M.K., Shin, B.A., and Choi, S.Y. (2011). High cleavage efficiency of a 2A peptide derived from porcine teschovirus-1 in human cell lines, zebrafish and mice. PLoS One 6, e18556.

Kramer, A., Yang, F.C., Snodgrass, P., Li, X., Scammell, T.E., Davis, F.C., and Weitz, C.J. (2001). Regulation of daily locomotor activity and sleep by hypothalamic EGF receptor signaling. Science 294, 2511–2515.

Kushikata, T., Fang, J., Chen, Z., Wang, Y., and Krueger, J.M. (1998). Epidermal growth factor enhances spontaneous sleep in rabbits. Am J Physiol 275, R509–514.

Lee, D.A., Andreev, A., Truong, T.V., Chen, A., Hill, A.J., Oikonomou, G., Pham, U., Hong, Y.K., Tran, S., Glass, L., et al. (2017). Genetic and neuronal regulation of sleep by neuropeptide VF. Elife 6.

Lee, D.A., Liu, J., Hong, Y., Hou, S., Hill, A.J., Lane, J.M., Wang, H., Oikonomou, G., Pham, U., Engle, J., et al. (2019). Evolutionarily Conserved Regulation of Sleep by Epidermal Growth Factor Receptor Signaling. Science Advances 5.

Lenz, O., Xiong, J., Nelson, M.D., Raizen, D.M., and Williams, J.A. (2015). FMRFamide signaling promotes stress-induced sleep in Drosophila. Brain Behav Immun 47, 141–148.

Lillesaar, C., Stigloher, C., Tannhauser, B., Wullimann, M.F., and Bally-Cuif, L. (2009). Axonal projections originating from raphe serotonergic neurons in the developing and adult zebrafish, Danio rerio, using transgenics to visualize raphe-specific pet1 expression. J Comp Neurol 512, 158–182.

Liu, D., and Dan, Y. (2019). A Motor Theory of Sleep-Wake Control: Arousal-Action Circuit. Annu Rev Neurosci 42, 27–46.

Liu, Q., Guan, X.M., Martin, W.J., McDonald, T.P., Clements, M.K., Jiang, Q., Zeng, Z., Jacobson, M., Williams, D.L., Jr., Yu, H., et al. (2001). Identification and characterization of novel mammalian neuropeptide FF-like peptides that attenuate morphine-induced antinociception. J Biol Chem 276, 36961–36969.

Ma, Y.J., Hill, D.F., Junier, M.P., Costa, M.E., Felder, S.E., and Ojeda, S.R. (1994). Expression of epidermal growth factor receptor changes in the hypothalamus during the onset of female puberty. Mol Cell Neurosci 5, 246–262.

Ma, Y.J., Junier, M.P., Costa, M.E., and Ojeda, S.R. (1992). Transforming growth factor-alpha gene expression in the hypothalamus is developmentally regulated and linked to sexual maturation. Neuron 9, 657–670.

Madelaine, R., Lovett-Barron, M., Halluin, C., Andalman, A.S., Liang, J., Skariah, G.M., Leung, L.C., Burns, V.M., and Mourrain, P. (2017). The hypothalamic NPVF circuit modulates ventral raphe activity during nociception. Sci Rep 7, 41528.

Mathias, J.R., Zhang, Z., Saxena, M.T., and Mumm, J.S. (2014). Enhanced cell-specific ablation in zebrafish using a triple mutant of Escherichia coli nitroreductase. Zebrafish 11, 85–97.

Mieda, M., Williams, S.C., Richardson, J.A., Tanaka, K., and Yanagisawa, M. (2006). The dorsomedial hypothalamic nucleus as a putative food-entrainable circadian pacemaker. Proc Natl Acad Sci U S A 103, 12150–12155.

Mikhail, C., Vaucher, A., Jimenez, S., and Tafti, M. (2017). ERK signaling pathway regulates sleep duration through activity-induced gene expression during wakefulness. Sci Signal 10.

Nagy, S., Tramm, N., Sanders, J., Iwanir, S., Shirley, I.A., Levine, E., and Biron, D. (2014). Homeostasis in C. elegans sleep is characterized by two behaviorally and genetically distinct mechanisms. Elife 3, e04380.

Nath, R.D., Chow, E.S., Wang, H., Schwarz, E.M., and Sternberg, P.W. (2016). C. elegans Stress-Induced Sleep Emerges from the Collective Action of Multiple Neuropeptides. Curr Biol 26, 2446–2455.

Nelson, M.D., Lee, K.H., Churgin, M.A., Hill, A.J., Van Buskirk, C., Fang-Yen, C., and Raizen, D.M. (2014). FMRFamide-like FLP-13 neuropeptides promote quiescence following heat stress in Caenorhabditis elegans. Curr Biol 24, 2406–2410.

Oikonomou, G., Altermatt, M., Zhang, R.W., Coughlin, G.M., Montz, C., Gradinaru, V., and Prober, D.A. (2019). The Serotonergic Raphe Promote Sleep in Zebrafish and Mice. Neuron 103, 686–701 e688.

Oikonomou, G., and Prober, D.A. (2017). Attacking sleep from a new angle: contributions from zebrafish. Curr Opin Neurobiol 44, 80–88.

Pollak Dorocic, I., Furth, D., Xuan, Y., Johansson, Y., Pozzi, L., Silberberg, G., Carlen, M., and Meletis, K. (2014). A whole-brain atlas of inputs to serotonergic neurons of the dorsal and median raphe nuclei. Neuron 83, 663–678.

Qian, Y., Cao, Y., Deng, B., Yang, G., Li, J., Xu, R., Zhang, D., Huang, J., and Rao, Y. (2017). Sleep homeostasis regulated by 5HT2b receptor in a small subset of neurons in the dorsal fan-shaped body of drosophila. Elife 6.

Ren, J., Friedmann, D., Xiong, J., Liu, C.D., Ferguson, B.R., Weerakkody, T., DeLoach, K.E., Ran, C., Pun, A., Sun, Y., et al. (2018). Anatomically Defined and Functionally Distinct Dorsal Raphe Serotonin Sub-systems. Cell 175, 472–487 e420.

Ren, J., Isakova, A., Friedmann, D., Zeng, J., Grutzner, S.M., Pun, A., Zhao, G.Q., Kolluru, S.S., Wang, R., Lin, R., et al. (2019). Single-cell transcriptomes and whole-brain projections of serotonin neurons in the mouse dorsal and median raphe nuclei. Elife 8.

Saper, C.B., and Fuller, P.M. (2017). Wake-sleep circuitry: an overview. Curr Opin Neurobiol 44, 186–192.

Scammell, T.E., Arrigoni, E., and Lipton, J.O. (2017). Neural Circuitry of Wakefulness and Sleep. Neuron 93, 747–765.

Schneider, C.A., Rasband, W.S., and Eliceiri, K.W. (2012). NIH Image to ImageJ: 25 years of image analysis. Nat Methods 9, 671–675.

Shang, Y., Donelson, N.C., Vecsey, C.G., Guo, F., Rosbash, M., and Griffith, L.C. (2013). Short neuropeptide F is a sleep-promoting inhibitory modulator. Neuron 80, 171–183.

Singh, C., Oikonomou, G., and Prober, D.A. (2015). Norepinephrine is required to promote wakefulness and for hypocretin-induced arousal in zebrafish. Elife 4.

Tabor, K.M., Bergeron, S.A., Horstick, E.J., Jordan, D.C., Aho, V., Porkka-Heiskanen, T., Haspel, G., and Burgess, H.A. (2014). Direct activation of the Mauthner cell by electric field pulses drives ultrarapid escape responses. J Neurophysiol 112, 834–844.

Turek, M., Besseling, J., Spies, J.P., Konig, S., and Bringmann, H. (2016). Sleep-active neuron specification and sleep induction require FLP-11 neuropeptides to systemically induce sleep. Elife 5.

Ubuka, T., Morgan, K., Pawson, A.J., Osugi, T., Chowdhury, V.S., Minakata, H., Tsutsui, K., Millar, R.P., and Bentley, G.E. (2009). Identification of human GnIH homologs, RFRP-1 and RFRP-3, and the cognate receptor, GPR147 in the human hypothalamic pituitary axis. PLoS One 4, e8400.

Urasaki, A., Morvan, G., and Kawakami, K. (2006). Functional dissection of the Tol2 transposable element identified the minimal cis-sequence and a highly repetitive sequence in the subterminal region essential for transposition. Genetics 174, 639–649.

Ursin, R. (2008). Changing concepts on the role of serotonin in the regulation of sleep and waking. In Serotonin and Sleep: Molecular, Functional and Clinical Aspects J.M. Monti, S.R. Pandi-Perumal, B.L. Jacobs, and D.J. Nutt, eds. (Switzerland: Birkhäuser Basel).

Van Buskirk, C., and Sternberg, P.W. (2007). Epidermal growth factor signaling induces behavioral quiescence in Caenorhabditis elegans. Nat Neurosci 10, 1300–1307.

van den Pol, A.N. (2012). Neuropeptide transmission in brain circuits. Neuron 76, 98–115.

Venner, A., Broadhurst, R.Y., Sohn, L.T., Todd, W.D., and Fuller, P.M. (2019). Selective activation of serotoninergic dorsal raphe neurons facilitates sleep through anxiolysis. Sleep.

Weber, F., and Dan, Y. (2016). Circuit-based interrogation of sleep control. Nature 538, 51–59.

Weissbourd, B., Ren, J., DeLoach, K.E., Guenthner, C.J., Miyamichi, K., and Luo, L. (2014). Presynaptic partners of dorsal raphe serotonergic and GABAergic neurons. Neuron 83, 645–662.

Westerfield, M. (1993). The zebrafish book : a guide for the laboratory use of zebrafish (Brachydanio rerio) (Eugene, OR: M. Westerfield).

Yelin-Bekerman, L., Elbaz, I., Diber, A., Dahary, D., Gibbs-Bar, L., Alon, S., Lerer-Goldshtein, T., and Appelbaum, L. (2015). Hypocretin neuron-specific transcriptome profiling identifies the sleep modulator Kcnh4a. Elife 4.

Yuan, Q., Joiner, W.J., and Sehgal, A. (2006). A sleep-promoting role for the Drosophila serotonin receptor 1A. Curr Biol 16, 1051–1062.

